# Ambroxol Hydrochloride Inhibits the Interaction between Severe Acute Respiratory Syndrome Coronavirus 2 Spike Protein’s Receptor Binding Domain and Recombinant Human ACE2

**DOI:** 10.1101/2020.09.13.295691

**Authors:** Omonike A. Olaleye, Manvir Kaur, Collins C. Onyenaka

## Abstract

Severe Acute Respiratory Syndrome Coronavirus 2 (SARS-CoV-2), the causative agent of coronavirus disease 2019 (COVID-19), enters the host cells through two main pathways, both involving key interactions between viral envelope-anchored spike glycoprotein of the novel coronavirus and the host receptor, angiotensin-converting enzyme 2 (ACE2). To date, SARS-CoV-2 has infected up to 26 million people worldwide; yet, there is no clinically approved drug or vaccine available. Therefore, a rapid and coordinated effort to re-purpose clinically approved drugs that prevent or disrupt these critical entry pathways of SARS-CoV-2 spike glycoprotein interaction with human ACE2, could potentially accelerate the identification and clinical advancement of prophylactic and/or treatment options against COVID-19, thus providing possible countermeasures against viral entry, pathogenesis and survival. Herein, we discovered that Ambroxol hydrochloride (AMB), and its progenitor, Bromhexine hydrochloride (BHH), both clinically approved drugs are potent effective modulators of the key interaction between the receptor binding domain (RBD) of SARS-CoV-2 spike protein and human ACE2. We also found that both compounds inhibited SARS-CoV-2 infection-induced cytopathic effect at micromolar concentrations. Therefore, in addition to the known TMPRSS2 activity of BHH; we report for the first time that the BHH and AMB pharmacophore has the capacity to target and modulate yet another key protein-protein interaction essential for the two known SARS-CoV-2 entry pathways into host cells. Altogether, the potent efficacy, excellent safety and pharmacologic profile of both drugs along with their affordability and availability, makes them promising candidates for drug repurposing as possible prophylactic and/or treatment options against SARS-CoV-2 infection.

## Introduction

The coronavirus disease 2019 (COVID-19) caused by severe acute respiratory syndrome coronavirus 2 (SARS-CoV-2) has emerged as a global pandemic and infected over 26 million individuals, with characteristics ranging from asymptomatic, mild, moderate to severe symptoms, with some people experiencing diverse multi-organ pathologies, substantial morbidity, and mortality^1–4^. Presently, there is no Food and Drug Administration (FDA) approved drug or vaccine available for treatment of COVID-19^5^. Although, there are several clinical studies investigating the possibility of repurposing existing clinically approved drugs as treatment options for COVID-19^5-9^; there still remains an urgent call for the rapid development of antiviral drugs for the prevention and/or treatment of COVID-19.

SARS-CoV-2 enters the host cells through two main pathways, both involving key interactions between viral envelope-anchored spike (S) glycoprotein of the novel coronavirus and the host receptor, angiotensin-converting enzyme 2 (ACE2), a membrane-bound metalloprotease^10–13^. The first pathway involves receptor mediated endocytosis process; while the second pathway, involves cell fusion also consisting of host receptor recognition and attachment of surface unit S1 to the peptidase domain of ACE2. Here, SARS-CoV-2 S protein is primed at the S1/S2 and the S2’ site by a plasma membrane-associated type II transmembrane serine protease (TMPRSS2), which triggers fusion of viral and cellular membranes, an essential step for release of the viral contents into the host cell cytosol^14,15^. X-ray crystallography and structural studies of human ACE2 revealed two domains; the N-terminal zinc metallopeptidase domain (MPD), which binds to the viral receptor binding domain (RBD) within the S glycoprotein, and a C terminal “collectrin-like” domain^16–18^. The crucial interaction between the MPD of human ACE2 and RBD of SARS-CoV-2 S glycoprotein has been characterized as an initial and critical step in viral infection^10-13^. Therefore, repurposing clinically approved drugs that could potentially prevent or disrupt this key interaction between SARS-CoV-2 S glycoprotein and human ACE2, could accelerate the clinical advancement of prophylactic and/or treatment options against COVID-19, thus providing possible countermeasures against viral entry, pathogenesis and survival.

Ambroxol hydrochloride ((AMB) 4-[(2-amino-3,5-dibromophenyl) methylamino]cyclohexan-1-ol; hydrochloride)^19^, belonging to the benzylamine structural class, is a demethylated active metabolite of Bromhexine hydrochloride (BHH) ^20^. Both AMB and its progenitor, BHH are widely prescribed drugs used to treat respiratory tract infections and disorders ^21–24^, clinically indicated for their secretolytic activity for treatment of acute and chronic bronchopulmonary diseases associated with abnormal mucus secretion and impaired mucus transport^20,21,24^. Traditionally, AMB and BHH have been available, affordable and used as over the counter drugs, with no significant adverse effects^24,25^. Furthermore, AMB and BHH have been extensively investigated in translational studies because of their multiple activities including mucociliary clearance activity, mucokinetic properties, stimulation of surfactant production, anti-inflammatory and antioxidative actions, and the local anesthetic effect ^22,23,26–28^. More recent studies have shown that AMB and BHH both induce cellular autophagic-lysosome pathway ^29–31^, critical processes in the host defense machinery against viral infections^32^. Additional studies have also shown the involvement of AMB in the modulation of homeostasis of ions such as, sodium, hydrogen, and calcium^33^. Moreover, AMB gained renewed interest as a potential drug for the clinical development of therapeutics for neurodegenerative diseases ^34^, due to its potential to act as a chaperone, pH-dependent, mixed-type inhibitor of glucocerebrosidase (GCase), and its involvement in mechanisms for mitochondria, lysosomal biogenesis, and secretory pathway ^31, 33, 34^. Other than the plethora of molecular impact and physiologic functions of the AMB pharmacophore mentioned above, reports have also shown that AMB could inhibit viruses that cause influenza and rhinovirus infections^35,36^. In addition, AMB’s progenitor BHH is an established potent inhibitor of TMPRSS2^37^, one of the key proteases for viral fusion into host cells. BHH’s activity against TMPRSS2 and lung protective properties makes it an attractive drug for the prevention and/or treatment of coronavirus infections ^38,39^. To date, COVID-19 clinical trials investigating the impact of BHH (and/or in combination with other drugs) on SARS-CoV-2 infection are ongoing or completed in some countries [NCT04273763; NCT04340349; NCT04355026; NCT04424134; NCT04405999 and IRCT202003117046797N4]. However, the exact mechanism of action of the BHH and AMB pharmacophore against SARS-CoV-2 infection needs to be elucidated.

In this study, we evaluated the effects of AMB, and its progenitor, BHH, on the interaction between recombinant human ACE2 (rhACE2) and the RBD of the S protein of SARS-CoV-2. We also determined the effect of both compounds on SARS-CoV-2 infection-induced cytopathic effect (CPE) *in vitro* and assessed their cytotoxicity, in comparison to other clinically approved drugs. Herein, we discovered that AMB and BHH, effectively modulated the rhACE2’s interaction with Spike (RBD) protein in the micromolar range. We also found that both compounds inhibited SARS-CoV-2 infection-induced CPE at certain concentrations. Therefore, in addition to the known TMPRSS2 activity of BHH^37^; we report for the first time that AMB and BHH pharmacophore have the capacity to target and modulate yet another key protein-protein interaction essential for the two known SARS-CoV-2 entry pathways in to human cells. Altogether, the potent efficacy, excellent safety and pharmacologic profile of both drugs along with their affordability and availability, makes them promising candidates for drug repurposing as possible prophylactic and/or treatment options against SARS-CoV-2 infection.

## MATERIALS AND METHODS

### MATERIALS

#### Cell Growth Conditions and Medium

African Green Monkey Kidney Vero E6 cells (ATCC# CRL-1586, American Tissue Culture Type) were maintained using medium purchased from Gibco (modified eagle’s medium (MEM) Gibco (#11095); 10% fetal bovine serum (HI FBS) Gibco (#14000); Penicillin/Streptomycin (PS) Gibco (#15140); 10U/mL penicillin and 10µg/mL streptomycin (only in assay media)). For the SARS-CoV-2 infection induced cytopathic effect (CPE) assay, cells were grown in MEM/10% HI FBS and harvested in MEM/1% PS/supplemented with 2% HI FBS. Cells were batch inoculated with SARS-CoV-2 USA_WA1/2020 (M.O.I. ∼ 0.002) which resulted in 5-10% cell viability 72 hours post infection.

#### Compounds and Preparation of Stock Solutions

We prepared 10mM stocks solutions of the inhibitors in Dimethyl sulfoxide (DMSO; D8418-Lot#SHBL5613) purchased from Sigma Aldrich. Ambroxol Hydrochloride (AMB; A9797 - Lot # BCCB1637), and Bromhexine Hydrochloride (BHH; 17343 - Lot # BCBJ8156V) were also purchased from Sigma Aldrich. Both compound samples were serially diluted 2-fold in DMSO nine times and screened in duplicates for the SARS-CoV-2 infection induced cytopathic effect assay. The reference compounds used for the CPE and cytotoxicity assays were made available by SRI. Assay Ready Plates (ARPs; Corning 3764BC) pre-drugged with test compounds (90 nL sample in 100% DMSO per well dispensed using a Labcyte (ECHO 550) were prepared in the Biosafety Level-2 (BSL-2) laboratory by adding 5µL assay media to each well.

#### Method for Measuring Antiviral Effect of AMB and BHH

Southern Research Institute (SRI), located in Birmingham, Alabama performed the SARS-CoV-2 infection induced cytopathic effect (CPE) assay and cytotoxicity assays through a sub-contract from Texas Southern University, Houston, Texas. The CPE reduction assay was conducted using a high throughput-screening (HTS) format as previously described ^40,41^. Specifically, Vero E6 cells selected for expression of the SARS-CoV-2 receptor (ACE2; angiotensin-converting enzyme 2) were used for the CPE assay. Cells were grown in MEM/10% HI FBS supplemented and harvested in MEM/1% PS/ supplemented with 2% HI FBS. Cells were batch inoculated with SARS-CoV-2 (M.O.I. ∼ 0.002) which resulted in 5% cell viability 72 hours post infection. Compound samples were serially diluted 2-fold in DMSO nine times and screened in duplicates. Assay Ready Plates (ARPs; Corning 3764 BC black-walled, clear bottom plates) pre-drugged with test compounds (90 nL sample in 100% DMSO per well dispensed using a Labcyte (ECHO 550) were prepared in the BSL-2 lab by adding 5µL assay media to each well. The plates were passed into the BSL-3 facility where a 25µL aliquot of virus inoculated cells (4000 Vero E6 cells/well) was added to each well in columns 3-22. The wells in columns 23-24 contained virus infected cells only (no compound treatment). Prior to virus infection, a 25µL aliquot of cells was added to columns 1-2 of each plate for the cell only (no virus) controls. After incubating plates at 37°C/5%CO_2_ and 90% humidity for 72 hours, 30µL of Cell Titer-Glo (Promega) was added to each well. Luminescence was read using a Perkin Elmer Envision or BMG CLARIOstar plate reader following incubation at room temperature for 10 minutes to measure cell viability. Raw data from each test well was normalized to the average (Avg) signal of non-infected cells (Avg Cells; 100% inhibition) and virus infected cells only (Avg Virus; 0% inhibition) to calculate % inhibition of CPE using the following formula: % inhibition = 100*(Test Cmpd - Avg Virus)/(Avg Cells – Avg Virus). The SARS CPE assay was conducted in BSL-3 containment with plates being sealed with a clear cover and surface decontaminated prior to luminescence reading.

#### Determination of Cytotoxic Effect of AMB and BHH

The cytotoxicity of AMB and BHH were assessed in a BSL-2 counter screen using the Cell Titer-Glo Luminescent Cell Viability Assay as previously described^41^. Briefly, host cells in media were added in 25µL aliquots (4000 cells/well) to each well of assay ready plates prepared with test compounds as above. Cells only (100% viability) and cells treated with hyamine at 100µM final concentration (0% viability) serve as the high and low signal controls, respectively, for cytotoxic effect in the assay. DMSO was maintained at a constant concentration for all wells (0.3%) as dictated by the dilution factor of stock test compound concentrations. After incubating plates at 37°C/5%CO2 and 90% humidity for 72 hours, 30µl CellTiter Glo (CTG) (G7573, Promega) was added to each well. Luminescence was read using a BMG CLARIOstar plate reader following incubation at room temperature for 10 minutes to measure cell viability.

#### SARS-CoV-2 Spike (RBD) Protein - ACE2 Interaction Assay

We purchased the SARS-CoV-2 Spike - ACE2 binding assay kits (Cat # CoV-SACE2-1, Lot# 062320 7066 and Lot# 081120 7066) from RayBiotech (Norcross, GA) and adapted the manufacturer’s protocol^42^ to determine the effect of AMB and BHH on the interaction between SARS-CoV-2 Spike (RBD) Protein and recombinant human ACE2. The *in vitro* enzyme-linked immunoabsorbent assay (ELISA) was performed in a transparent flat-bottom 96-well plate. We prepared 10mM stock solutions of the compounds in Dimethyl sulfoxide (DMSO), with serially diluted the compounds in DMSO as follows: 100, 50, 10, 5, 1, 0.5, and 0.1 µM for AMB and BHH. All experiments were performed in triplicates. Each plate contained positive controls (1% DMSO) and blank controls with no ACE2. Specifically, 1 µL of serially diluted compounds were incubated with recombinant SARS-CoV-2 Spike receptor binding domain (RBD) protein, pre-coated on the 96 well plates in 49 µL of 1X assay diluent buffer for about 30 mins, at room temperature (22°C) with shaking at 180 rpm. Then, we added 50 µL of ACE2 protein in 1X assay diluent buffer into the 96 well plate, and incubated for 2.5 hrs at room temperature (22°C) with shaking at 180 rpm. Thereafter, the solution was discarded and the plate was washed consecutively four times with 300 µL 1X wash buffer, followed by the addition of the detection antibody (anti-ACE2 goat antibody). The reaction was allowed to go on for 1 hr at room temperature (22°C) with shaking at 180rpm. Then, the solution was discarded and the wash step was repeated as described above. Next, the HRP-conjugated anti-goat IgG was added to each well, and the reaction plate was further incubated for 1 hr at room temperature (22°C) with shaking at 180rpm. Again, the solution was discarded and the wash step was repeated as described above. Then, 100µL of 3,3’,5,5’-tetramethylbenzidine (TMB) one-step substrate was added to each well, and reaction mixtures were incubated in the dark at room temperature (22°C) with shaking at 180rpm for 30 mins. The reaction was stopped by the addition of 50 µL stop solution. The absorbance was read at 405 nm using a Beckman Coulter DTX880 multimode plate reader. The background hydrolysis was subtracted and the data was fitted to a special bell-shaped dose-response curve equation using GraphPad prism software 8.4.3.

## RESULTS

### Effects of AMB and BHH on Spike (RBD) Glycoprotein and rhACE2 Interaction

The interaction of human ACE2 and the receptor binding domain (RBD) of the SARS-CoV-2 S protein has been reported as a critical step in viral infection for both the endocytosis and non-endocytosis pathways of viral entry into host cells ^10-15, 43^. We evaluated the effect of AMB and its progenitor, BHH on the binding affinity of rhACE2 and RBD of SARS-CoV-2 S protein at concentrations ranging from 100 µM to 100 nM, using an adapted *in vitro* enzyme-linked immunoabsorbent assay (ELISA)^42^. We observed a unique dose-response curve for both compounds (using the special bell shape curve model). AMB displayed the highest inhibition of S (RBD)-rhACE2 protein interaction at lower micromolar concentrations (ranging from 100nM to 10µM); compared to higher concentrations of AMB from 50 µM (Figure 1B). Interestingly, we found that BHH inhibited the binding of SARS-CoV-2’s S (RBD) protein to rhACE2 receptor at lower concentrations ranging from 100 nM to 10 µM; but enhanced the interaction at higher concentrations of BHH from 50 µM (Figure 1A). Hence, the bell shaped model generated two IC_50_ values (IC_50_1_ and IC_50_2_) as shown in Table 4. Unlike BHH, we did not observe a stimulation or enhancement of binding of SARS-CoV-2’s Spike (RBD) protein to rhACE2 receptor for its metabolite – AMB, at the concentrations we tested. However, using the bell curve model, the graphpad software generated a second IC_50_2_ at 232 µM for AMB, greater than the highest concentration tested (100 µM). Moreover, the unconventional dose response curve observed in this protein interaction assay, could be intrinsic to the mode of inhibition or an indicator of additional binding site(s) and/or target(s), for the BHH and AMB, such as other sites on rhACE2 or the Spike (RBD) protein. To our knowledge, these finding is the first report to reveal that AMB inhibits and interferes with the binding between rhACE2 receptor and SARS-CoV-2 S (RBD) glycoprotein *in vitro*; while BHH inhibits this critical interaction at lower concentrations and enhances at higher concentrations (Table 4). These results suggest that AMB might be a promising lead for clinical development of novel SARS-CoV-2 entry inhibitors and potential COVID-19 therapeutics (Figure 2). Altogether, these results reveal a new pharmacologic mode of action and novel target for AMB and its progenitor, BHH.

**Figure 1.**
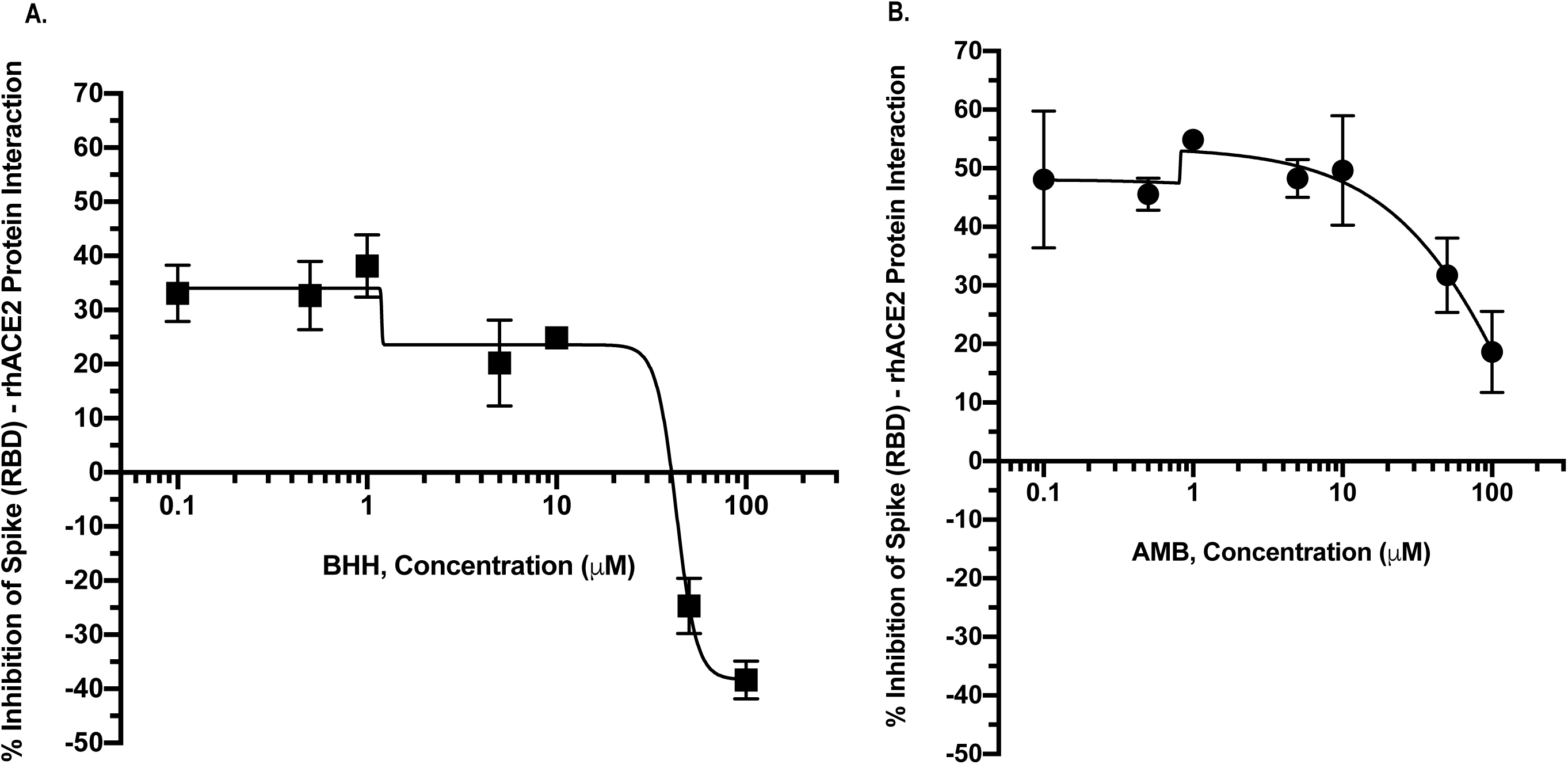
Effect of Bromhexine Hydrochloride (BHH) and Ambroxol Hydrochloride (AMB) on the interaction of rhACE2 with SARS-CoV-2 Spike (RBD) protein Interaction: A. BHH, and B. AMB.

**Figure 2.**
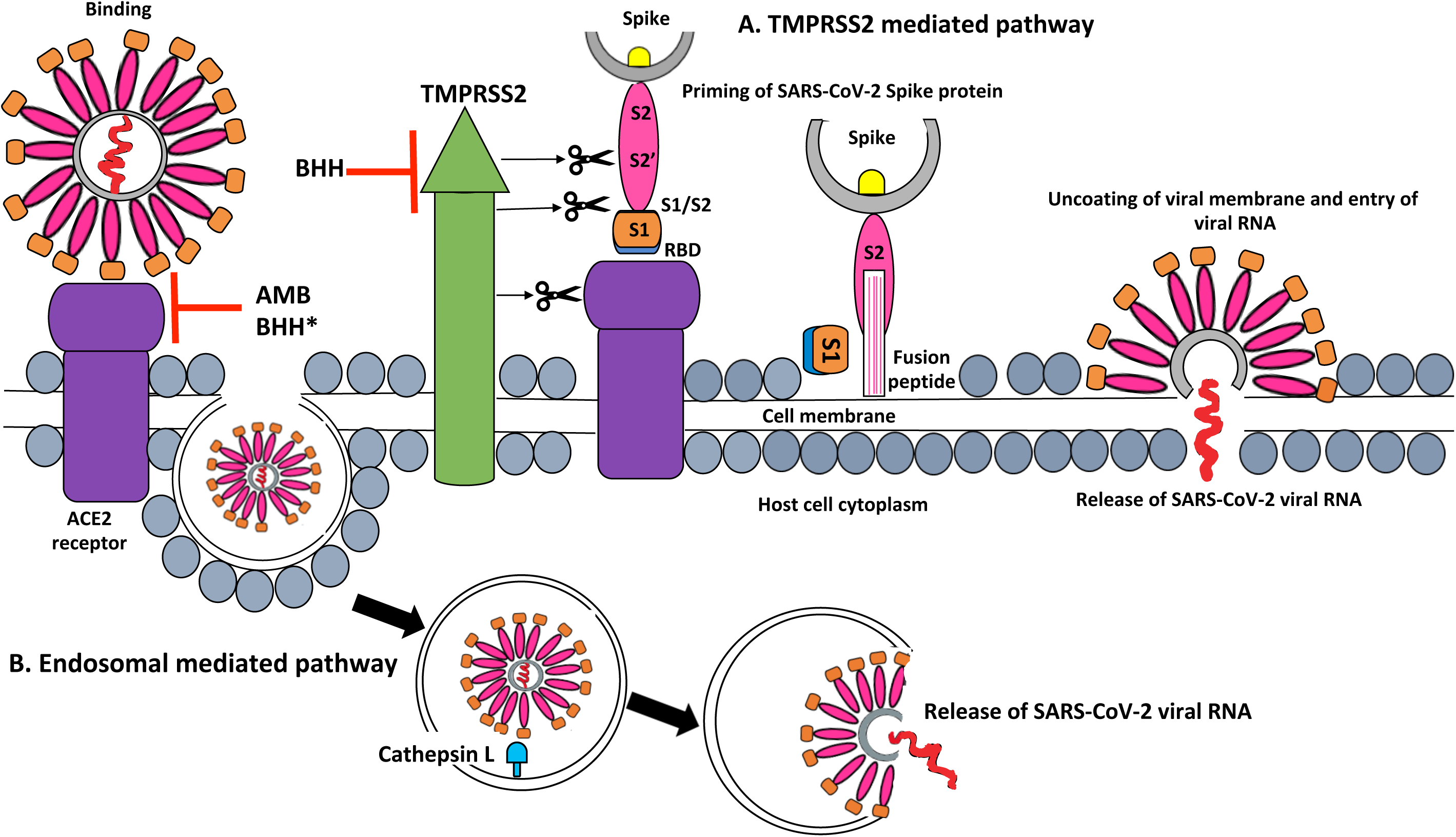
Schematic diagram depicting SARS-CoV-2 Spike (RBD) protein interaction with human ACE2 receptor at the cellular membrane, and inhibition of the two entry pathways of virus into host cells by Ambroxol Hydrochloride (AMB) and Bromhexine Hydrochloride (BHH) pharmacophore: A. TMPRSS2 mediated pathway, and B. Endosomal pathway activated by Cathepsin L. *At high concentrations, BHH was shown to also enhance the interaction between recombinant hACE2 and SARS-CoV-2 Spike (RBD) protein.

### Efficacy of Ambroxol Hydrochloride (AMB) and Bromhexine Hydrochloride (BHH) against SARS-CoV-2 infection induced Cytopathic Effect (CPE) in Vero E6 cells

The rapid identification and repurposing of clinically approved drugs with antiviral activity against SARS-CoV-2 infection could potentially accelerate their consideration as potential treatment options for COVID-19. Here in, we evaluated the *in vitro* antiviral activity of AMB, and its progenitor, BHH, using a standard luminescent-based high-throughput screening (HTS) platform^40,41^ for SARS-CoV-2 infection induced CPE in African Green Monkey Kidney Vero E6 cells. We found that, BHH inhibited SARS-CoV-2 infection induced CPE *in vitro* with 50% Inhibitory Concentration (IC_50_) value at about 21.72 µM; while AMB’s IC_50_ was greater than 30 µM, the highest concentration tested (Table 1). At this maximum concentration, AMB displayed 14.25% inhibition of SARS-CoV-2 induced CPE. Compared to its metabolite; BHH exhibited the highest maximum inhibition at about 91.08% inhibition at 30 µM (Table 1). Additional higher concentrations need to be tested in future studies to determine the actual IC_50_ value of AMB against SARS-CoV-2 infection induced CPE in Vero E6 cells. Furthermore, we compared the antiviral effects of AMB and BHH with five other known inhibitors of SARS-CoV-2 *in vitro*: Calpain Inhibitor IV, Chloroquine, Remdesivir, Hydroxychloroquine and Aloxistatin. We found that the IC_50_ for most of the reference compounds (Calpain Inhibitor IV (0.29µM), Chloroquine (3.56µM), Hydroxychloroquine (5.16µM), Remdesivir (8.54µM)) were lower than the IC_50S_ values for BHH and AMB. However, the IC_50_ of Aloxistatin (21.78 µM) was similar to that of BHH (21.72 µM) (Table 1 and 2). The IC_50_ values that we observed for the reference compounds are consistent with earlier reports ^44–47^. While writing this manuscript, we found that other cellular studies tested BHH and AMB at certain or single concentrations ^48–50^; however, this is the first study to our knowledge to conduct the IC_50_ evaluation for BHH and AMB against the novel SARS-CoV-2 infection induced CPE.

**Table 1.**
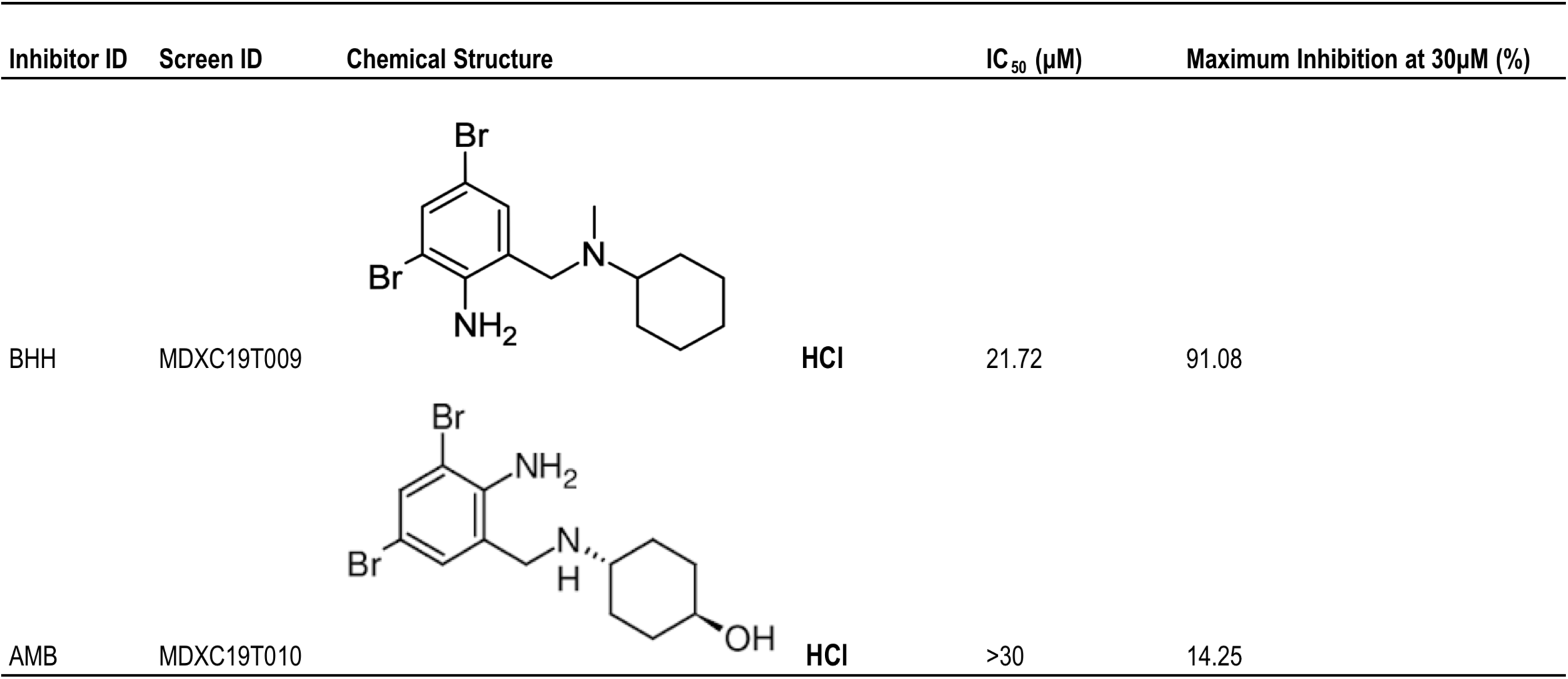
Chemical Structure and Activity of Ambroxol Hydrochloride (AMB) and Bromhexine Hydrochloride (BHH) against SARS-CoV-2 induced Cytopathic Effect (CPE) in Vero E6 Cells.

| Inhibitor ID | Screen ID | Chemical Structure |  | IC<sub>50</sub> (μM) | Maximum Inhibition at 30μM (%) |
| --- | --- | --- | --- | --- | --- |
| BHH | MDXC19T009 |  | HCl | 21.72 | 91.08 |
| AMB | MDXC19T010 |  | HCl | >30 | 14.25 |

**Table 2.**
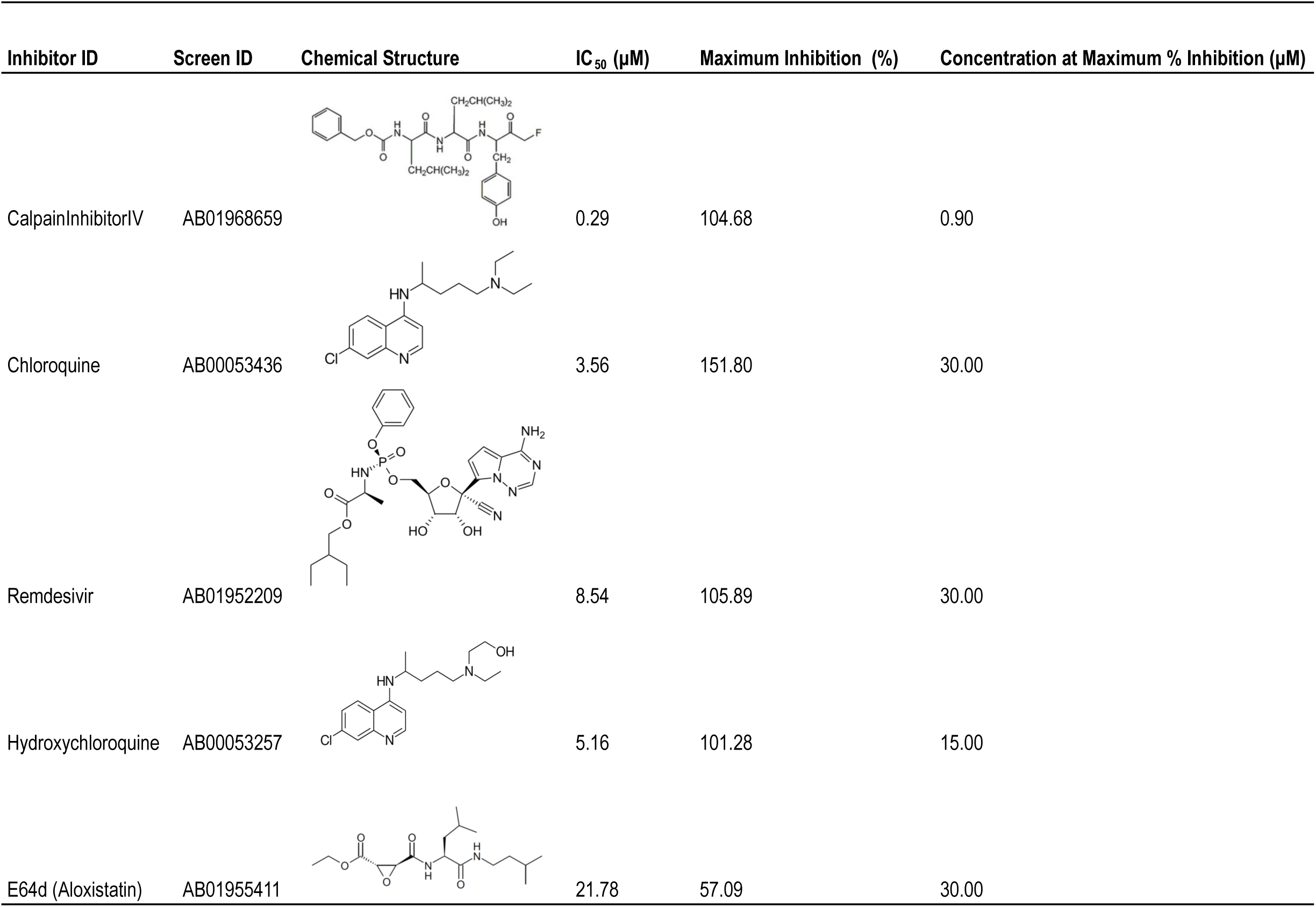
Chemical Structure and Activity of Reference Inhibitors against SARS-CoV-2 induced Cytopathic Effect (CPE) in Vero E6 Cells.

| Inhibitor ID | Screen ID | Chemical Structure | IC<sub>50</sub> (μM) | Maximum Inhibition (%) | Concentration at Maximum % Inhibition (μM) |
| --- | --- | --- | --- | --- | --- |
| CalpainInhibitorIV | AB01968659 |  | 0.29 | 104.68 | 0.90 |
| Chloroquine | AB00053436 |  | 3.56 | 151.80 | 30.00 |
| Remdesivir | AB01952209 |  | 8.54 | 105.89 | 30.00 |
| Hydroxychloroquine | AB00053257 |  | 5.16 | 101.28 | 15.00 |
| E64d (Aloxistatin) | AB01955411 |  | 21.78 | 57.09 | 30.00 |

### Cytotoxicity Effects of AMB and BHH in Vero E6 cells

Using a Cell Titer-Glo Luminescent Cell Viability Assay^41^, we determined the cytotoxicity of AMB and BHH. We also assessed the cytotoxic effects of the reference compounds in Vero E6 cells and observed that, the 50% cytotoxic concentration (CC_50_) of AMB and BHH were greater than 30 µM. In comparison to the other reference compounds tested, AMB and BHH displayed slightly higher percent maximum and minimum viability at the concentrations tested (Table 3). Between the two compounds, we observed a higher percent minimum viability for AMB (113.95%), compared to its progenitor, BHH (103.87%) at 30 µM (Table 3). These cytotoxicity results are consistent with the known clinical safety profiles of both compounds, with AMB showing better pharmacokinetic and safety profiles compared to BHH^51^.

**Table 3.** Cytotoxicity of Ambroxol Hydrochloride (AMB) and Bromhexine Hydrochloride (BHH) in Vero E6 Cells, in Comparison to Reference Inhibitors of SARS-CoV-2.

| Inhibitor ID | Cytotoxicity CC<sub>50</sub> (μM) | Minimum Viability (%) | Concentration at Minimum % Viability (μM) | Maximum Viability (%) | Concentration at Maximum % Viability (μM) |
| --- | --- | --- | --- | --- | --- |
| BHH | >30.00 | 103.87 | 30.00 | 133.17 | 0.12 |
| AMB | >30.00 | 113.95 | 30.00 | 124.37 | 1.88 |
| CalpainInhibitorIV | >7.17 | 97.74 | 7.17 | 113.22 | 0.45 |
| Chloroquine | >30.00 | 93.52 | 30.00 | 111.43 | 0.06 |
| Remdesivir | >30.00 | 101.07 | 0.120 | 109.76 | 15.00 |
| Hydroxychloroquine | >30.00 | 96.10 | 0.470 | 105.31 | 0.12 |
| E64d (Aloxistatin) | >30.00 | 97.45 | 30.000 | 104.15 | 0.47 |

**Table 4.** Activity of Ambroxol Hydrochloride (AMB) and Bromhexine Hydrochloride (BHH) against rhACE2 and SARS-CoV-2 Spike (RBD) protein Interaction.

| Estimated Relative IC<sub>50</sub> (μM) for Spike (RBD)-rhACE2 Protein Interaction Assay |  |  |
| --- | --- | --- |
| Inhibitor ID | IC<sub>50_1</sub> (μM) | IC<sub>50_2</sub> (μM) |
| BHH | 1.19 | 42.90 |
| AMB | 0.82 | 231.60 |

## DISCUSSION

The emergence of novel SARS-CoV-2 strains has imposed an urgent call to accelerate the repurposing and advancement of existing clinically safe and approved drugs as effective prophylactic and/or therapeutic agents for combating the COVID-19 pandemic. The crystal structure of full length human ACE2 revealed that the RBD on SARS-CoV-2 S1 binds directly to the metallopeptidase domain (MPD) of ACE2 receptor^10-13^. In this study, we explored the possibility of a clinically safe and approved drug, AMB and its progenitor, BHH as potential effectors of the interaction between SARS-CoV-2’s Spike protein receptor binding domain and recombinant human ACE2 receptor, a critical first step in the two significant pathways required for viral entry into host cells^10-15,43^ and initiation of pathogenesis (Figure 2). Using a sensitive ELISA^42^, we found that AMB and its progenitor, BHH modulates the interaction of rhACE2 and S (RBD) protein, with AMB being the most potent effector. Significant inhibition of the interaction between SARS-CoV-2’s S (RBD) protein and rhACE2 by AMB at nanomolar to micromolar concentrations provides strong evidence that this pharmacophore could serve as a promising lead for the discovery and clinical development of novel SARS-CoV-2 entry inhibitors and potential COVID-19 therapeutics. On the other hand, we found that BHH, enhances this key interaction between Spike (RBD) protein and rhACE2 at higher concentrations; while inhibiting at lower concentrations. To our knowledge, this is the first experimental study revealing this class of compounds as potent effectors of the binding of Spike (RBD) protein to rhACE2, a new pharmacologic mode of action for potentially modulating viral entry into host cells. *In vivo* pharmacodynamics and pharmacokinetic studies are warranted to further explore the physiologic relevance of the inhibitory activity of AMB, as well as the dual inhibitory and enhancement activities of BHH in the context of SARS-CoV-2 infection.

Our findings regarding the “paradoxical behavior” of BHH *in vitro* biochemical assay, are consistent with a recent report by Hörnich, B.F. et. al., that revealed that BHH enhanced SARS-CoV-2 S-mediated fusion in 293T cells at the concentrations tested^48^. Moreover, the unconventional dose-response curve that we observed in our interaction studies suggests that there may be more than one binding site for interaction of rhACE2 and Spike (RBD) protein for BHH and its metabolite-AMB, resulting in the potent inhibition of interaction at lower micromolar concentrations; compared to the impact at higher concentrations. This may also explain the enhancement of interaction by BHH at higher concentrations. Interestingly, a molecular dynamic study by Silva de Souza et al.^52^, revealed that two different regions within the RBD of SARS-CoV-2 interact differently with hACE2 in the presence of high salt concentrations (E1, is more hydrophobic; while E2 favors more polar interactions). Hence, suggesting that these differences may impact how effectors modulate these sites and affect the binding between RBD to hACE2. Future computational modeling and X-ray co-crystallization studies of the compounds with rhACE2 and Spike (RBD) may provide additional insight into the mode of inhibition (for AMB) and/or enhancement in the case of BHH. Besides, this will facilitate the rational design of novel drugs that could inhibit this interaction and in turn, prevent viral entry into human cells. In addition, synthesis of more potent inhibitors in the AMB-containing, benzylamine structural class and SAR studies could accelerate the discovery of novel anti-SARS-CoV-2 agents.

Furthermore, we investigated the effect of AMB and BHH on SARS-CoV-2 infection induced CPE *in vitro*, using a simple and rapid cellular high throughput screening assay^40,41^. Although, AMB was found to be the most potent against rhACE2-RBD interaction in the biochemical assay compared to BHH; however, in the cellular CPE assay, AMB had a higher IC_50_ than BHH for antiviral activity. Additionally, we observed that the IC_50_ values of the compounds in the cellular CPE assay were much higher than the IC_50s_ in the rhACE2-Spike(RBD) protein interaction assay, although the special bell curve model produced two IC_50s_ due to the mode of inhibition at lower concentrations versus higher concentrations. Therefore, *in vivo* pharmacokinetic and pharmacodynamics studies will be required to ascertain half maximal effective concentrations (EC_50_) for both drugs. Consistent with our study, a high throughput screen of clinically approved drugs by Tourte et. al. ^50^ also identified AMB as an inhibitor of SARS-CoV-2 infection *in vitro* at 10 µM. Our findings for percent inhibitions for AMB in the CPE assay in Vero E6 cells are also consistent with a recent study by Bradfute, S.B. et. al^49^., in which they reported that AMB could inhibit SARS-CoV-2 infection *in vitro* at high concentrations^49^. Our biochemical findings are most complementary to the report by Hornich, B.F. et. al., that observed inhibition of SARS-CoV-2-S-mediated fusion in the presence of 50 µM AMB^48^; but observed a dose dependent enhancement of SARS-CoV-2-S-mediated fusion by BHH at the concentrations tested^48^. Furthermore, a computational transcriptomics study by Napolitano, F. et al. ^53^, found that AMB induced an opposite transcriptional profile compared to that induced by SARS-CoV-2 infection *in vitro*, suggesting that it may be playing a role in counteracting viral infection^53^. Altogether, our findings provide a novel protein-protein interaction target and strong supporting evidence that may explain the mechanism of action for the AMB and BHH pharmacophore (Figure 2).

In addition, we conducted a comparative analysis of the dose-response curves of antiviral activity and cytotoxicity of AMB and BHH with five other known inhibitors of SARS-CoV-2 *in vitro*, which we refer to as “reference compounds.” We found that the IC_50_ range for BHH and AMB were similar to that of Aloxistatin (Table 1 and 2); but higher than the other reference compounds, namely, Chloroquine, Hydroxychloroquine, Remdesivir, and Calpain Inhibitor IV. On the other hand, when we assessed the cytotoxicity of AMB and BHH in Vero E6 cells, we observed that both compounds displayed higher percent maximum and minimum viability at the concentrations tested (Table 3) compared to the other reference compounds. Moreover, AMB had a slightly higher percent minimum viability compared to BHH, consistent with other reported safety studies for both compounds, that demonstrate that AMB has superior safety profile than BHH^51^. Thus, considering that AMB is safe and clinically approved, our findings that AMB significantly inhibited binding of rhACE2 receptor with SARS-CoV-2 Spike (RBD) protein and moderately inhibited SARS-CoV-2 infection induced CPE at the concentration tested, strongly supports the notion that AMB could be a safe drug option for the clinical advancement of counter measures against COVID-19. In addition, AMB and its progenitor – BHH, could be used as chemical probes to study the biology of host-pathogen interaction in the context of SARS-CoV-2 infections, particularly in the pre-clinical development of novel entry inhibitors. Our findings not only provides strong *in vitro* evidence and mechanism, for the proposed use of AMB as a therapeutic option for the treatment of COVID-19; but also sheds light on the relevance of this pharmacophore as potential prophylactic options for prevention of SARS-CoV-2 infection.

Both AMB and BHH have been used in the clinic for treatment of respiratory conditions and are extensively studied, because of their multiple pharmacologic effects and safety profile^20-24^. In addition to their impact on lung physiology and function with regards to mucociliary clearance, mucokinetic properties, and stimulation of surfactant production, they have also been shown to elicit anti-inflammatory, antioxidative and anesthetic effects^20-24^ Additional studies have reported that both compounds induce cellular autophagic-lysosome pathway ^29-31^. Reports have also shown the involvement of AMB in modulation of the homeostasis of ions such as sodium, hydrogen, and calcium^33, 51^. Additionally, AMB has gained attention clinically as a potential drug to repurpose for treatment of neurodegenerative diseases^31, 34^. Moreover, AMB is known to inhibit certain viruses *in vitro* and *in vivo*, with one proposed mechanism as that of preventing the release of RNA into the cytoplasm by increasing the endosomal pH^35,36^. This may suggest the possibility of additional mechanisms of action beyond viral entry pathways, that is, post-entry of virus into host cells. Given the ongoing COVID-19 pandemic and the lack of clinically approved vaccine or drugs; AMB and its progenitor, BHH could be possible drug leads, because of earlier reports of BHH’s inhibitory activity against TMPRSS2 at low micromolar concentrations^37^. However in a recent study, unlike Camostat, another known TMPRSS2 inhibitor; AMB and BHH did not reverse TMPRSS2-mediated enhancement of SARS-CoV-2 infection in 293Tcells^48^. Suggesting that AMB and BHH, may have additional modes/sites of action^48^. Hence, our finding that AMB and BHH is an effector of the RBD-rhACE2 interaction could shed more light on the mode of SARS-CoV-2 inhibition.

The clinical advancement of the AMB pharmacophore could potentially accelerate their consideration for repurposing as potential antiviral agents against COVID19, in non-human primate (NHP) models of SARS-CoV-2 infection, and in clinical trials. Ongoing COVID19 clinical trials [NCT04273763; NCT04340349; and NCT04355026; NCT04424134; NCT04405999] investigating the impact of BHH use prophylactically and/or for treatment in combination therapy for SARS-CoV-2 infection may provide more insight on the relevance of this pharmacophore in counteracting viral entry, pathogenesis and survival. To date, the only published clinical trials results of BHH, is that of a completed open-label randomized clinical trial conducted in Tabriz, North West of Iran ^54^. This small sample-size study, revealed that early administration of oral BHH decreases the intensive care unit transfer, intubation, and the mortality rate in patients with COVID-19^54^. Additional large scale and multi-country trials are needed to assess and ascertain the effect of BHH and/or AMB on clinical outcomes and mortality in COVID-19 patients. Moreover, previous studies by others have shown that both AMB and BHH could enhance the lung levels of certain antibiotics when used in combination^29,30^, suggesting that combination therapy with other known antimicrobials with antiviral activity against SARS-CoV-2 could be explored in COVID-19 clinical studies. Therefore, we propose the clinical evaluation and comparative analysis of the impact of AMB or BHH (or in combination with other available therapeutics) to reduce COVID19 morbidity and mortality.

Overall, the abundance of clinical evidence on the pharmacologic spectrum of activity of AMB, combined with its extensive clinical safety profile, affordability and availability, makes this an attractive and promising drug for targeting the binding of SARS-CoV-2 Spike (RBD) to rhACE2, an essential biologic interaction for viral entry into host cells^10-13^. The unique potential of AMB and/or BHH in modulating the two currently known viral entry pathways, makes this pharmacophore promising as possible prophylactic agents against SARS-CoV-2 infection. One of the main strengths of our study is the use of a rapid and sensitive ELISA, to identify and characterize two existing clinical drugs as novel inhibitors of the critical interaction between SARS-CoV-2 Spike (RBD) protein and rhACE2. However, our study has some limitations such as the use of Vero E6 cells that were selected for high expression of human ACE2 in the antiviral assay^40,41^. Besides, future studies exploring structural activity relationship (SAR) with derivatives of AMB, with a similar pharmacophore, improved efficacy and similar/better safety profile, will facilitate the discovery and pre-clinical development of new chemical molecules as potential options for COVID-19 prevention and/or treatment. SAR studies will also provide additional insight into the mode of action of this pharmacophore and aid in the rational design/discovery of new countermeasures against SARS-CoV-2 infection.

## CONCLUSION AND SIGNIFICANCE

Repurposing clinically approved drugs with novel mechanisms of action and/or multiple cellular targets could potentially disrupt viral pathogenesis, survival and/or prevent the viral entry and interaction with host receptor, ACE2. This approach to drug discovery could accelerate the clinical development of anti-COVID-19 prophylactics/treatments, thus providing countermeasures against COVID-19 and its impact on population health, healthcare systems, and the global economy^55^. Our novel findings of AMB and BHH as modulators of the binding of rhACE2 with SARS-CoV-2 Spike (RBD) protein provide new insights into the mode of action and additional molecular target(s) for AMB and its progenitor, BHH. Thus, validating this chemical class as promising leads for clinical development of novel SARS-CoV-2 entry inhibitors that could be used for potential COVID-19 prevention and/or treatment. Additionally, because these pharmacophore was shown earlier to have activity against TMPRSS2; and now we found that it inhibits rhACE2-RBD interaction, thus, probably targeting more than one protein. Therefore, this pharmacophore represents a favorable drug class that could possibly be studied as lead series in the development of countermeasures for limiting SARS-CoV-2 drug resistance in the future. Also its clinical efficacy as a secretolytic and anti-inflammatory agent makes it a promising drug for COVID-19 treatment due to lung injury. Therefore, we propose additional pharmacologic and clinical studies to further explore AMB and/or its derivatives as prophylaxis and/or treatment options in the toolbox for combating this novel coronavirus.

## Author Contributions

OAO conceived the study and performed biochemical experiments (rhACE2-Spike (RBD) protein interaction experiments); OAO and MK performed experimental design, data analysis and interpretations. OAO and MK wrote the manuscript. CCO created Figure 2a and 2b and helped to thoroughly review and edit the manuscript.

